## Supplementary figures and images for "Turnover of retroelements and satellite DNA drives centromere reorganization over short evolutionary timescales in Drosophila"

### Sup Fig1

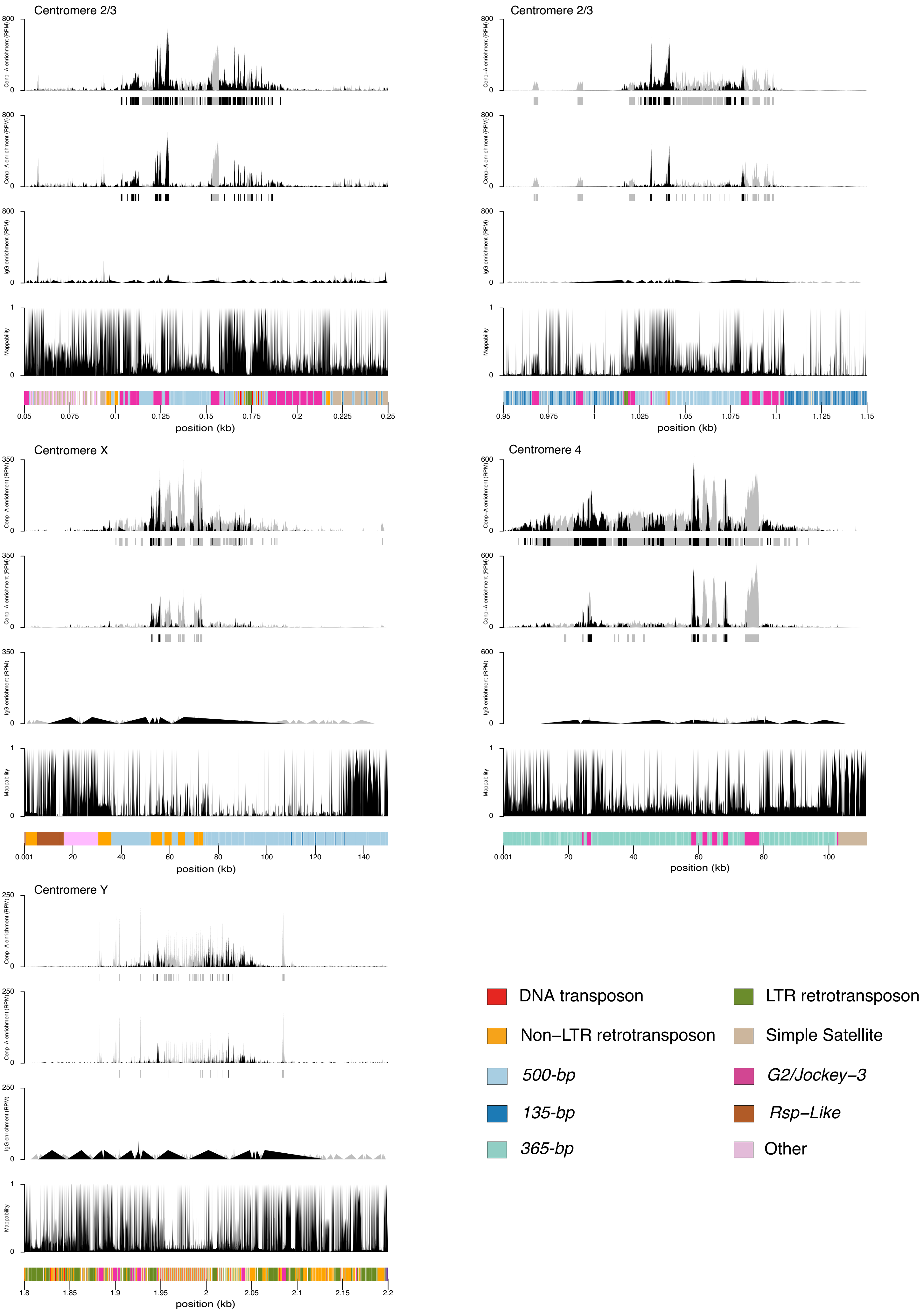

### Sup Fig2

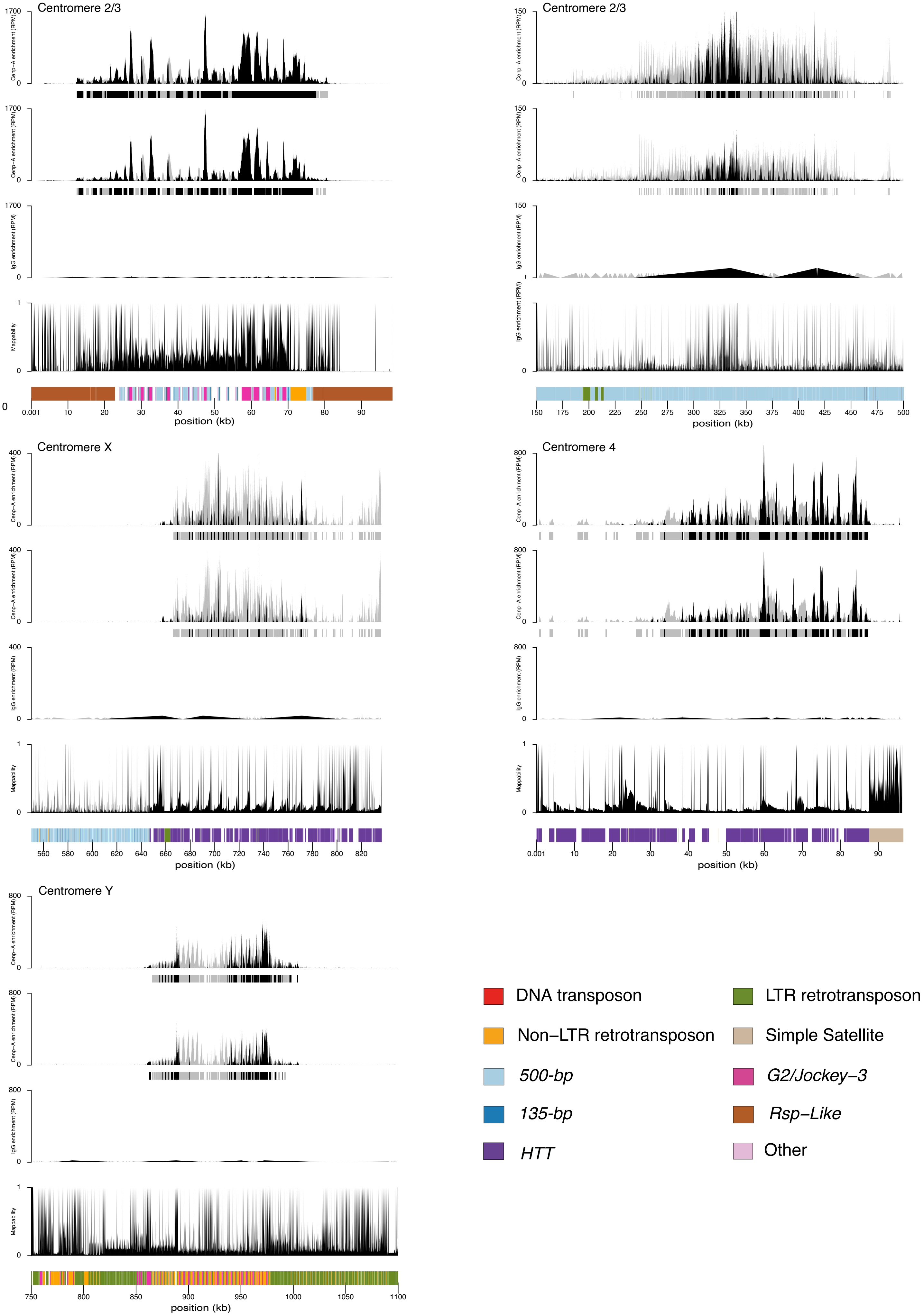

### Sup Fig3

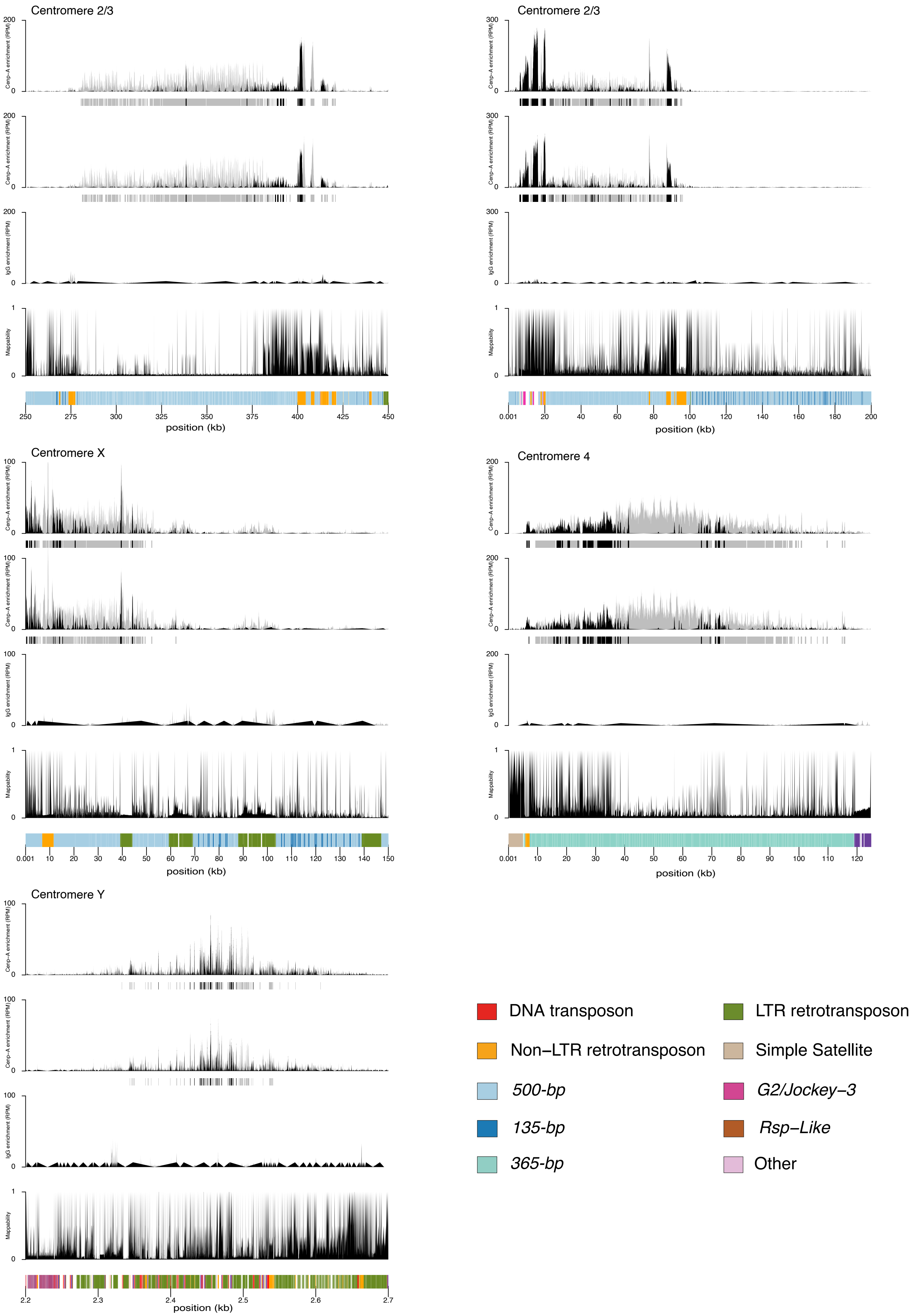

### Sup Fig4

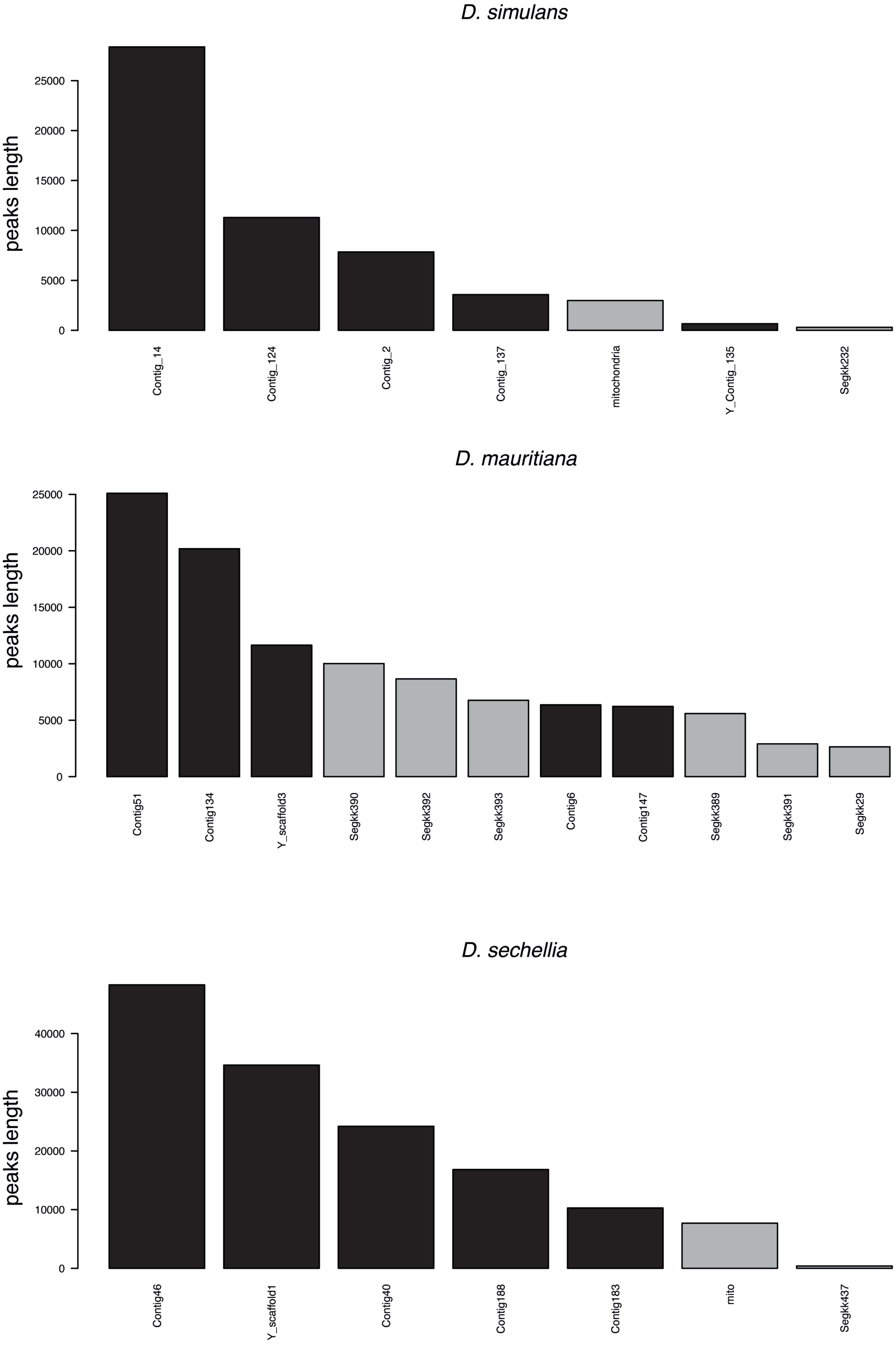

### Sup Fig5

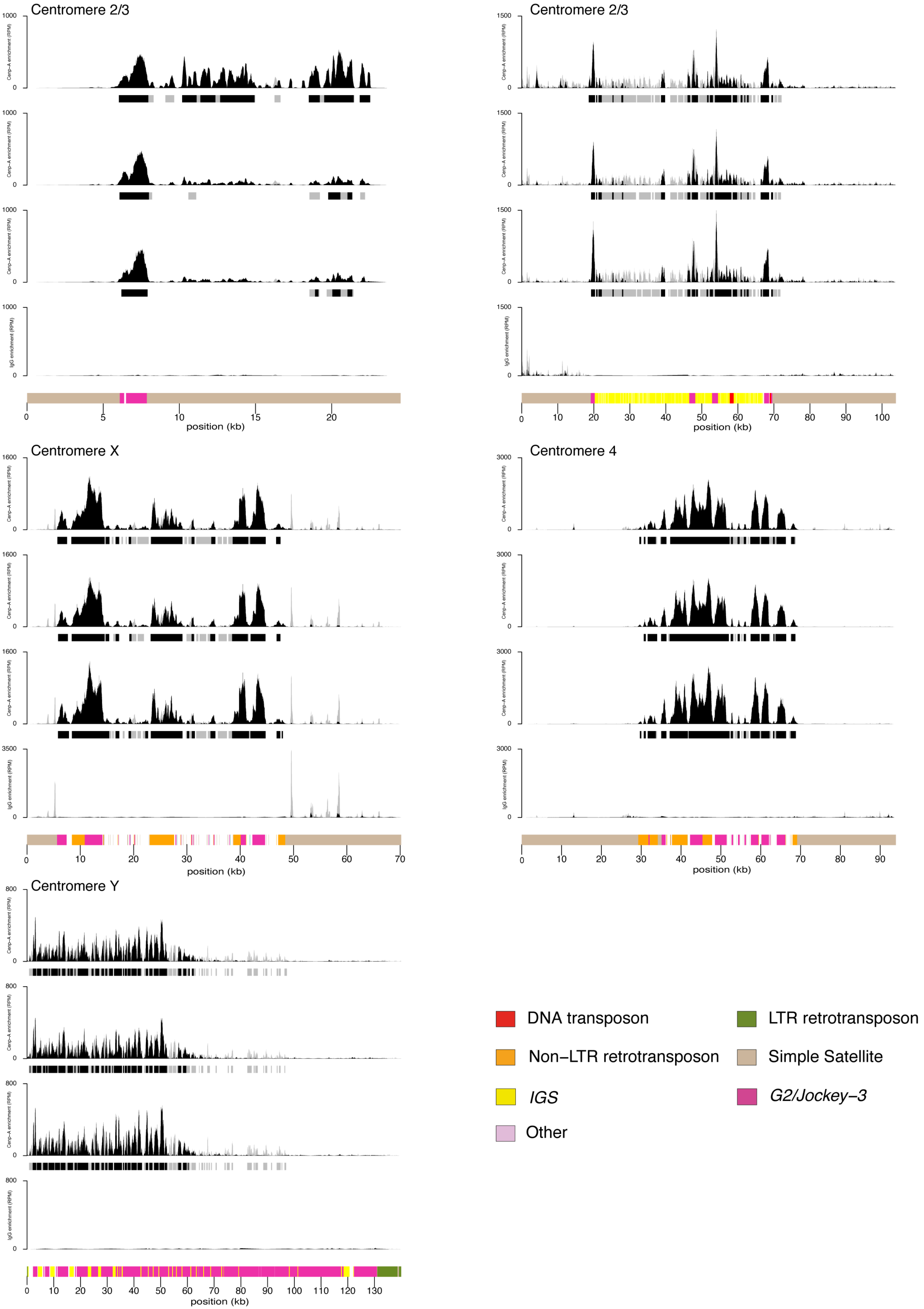

### Sup Fig6

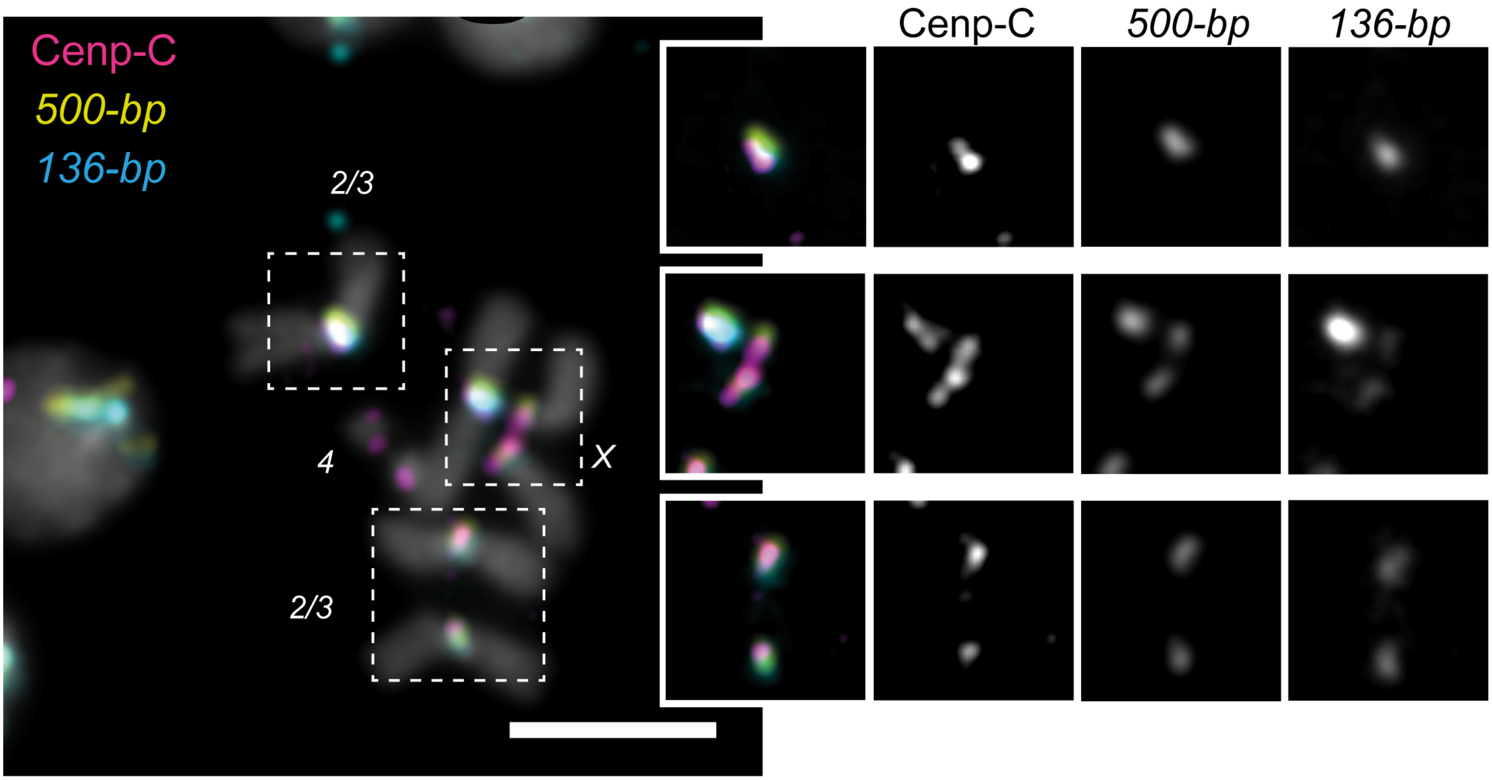

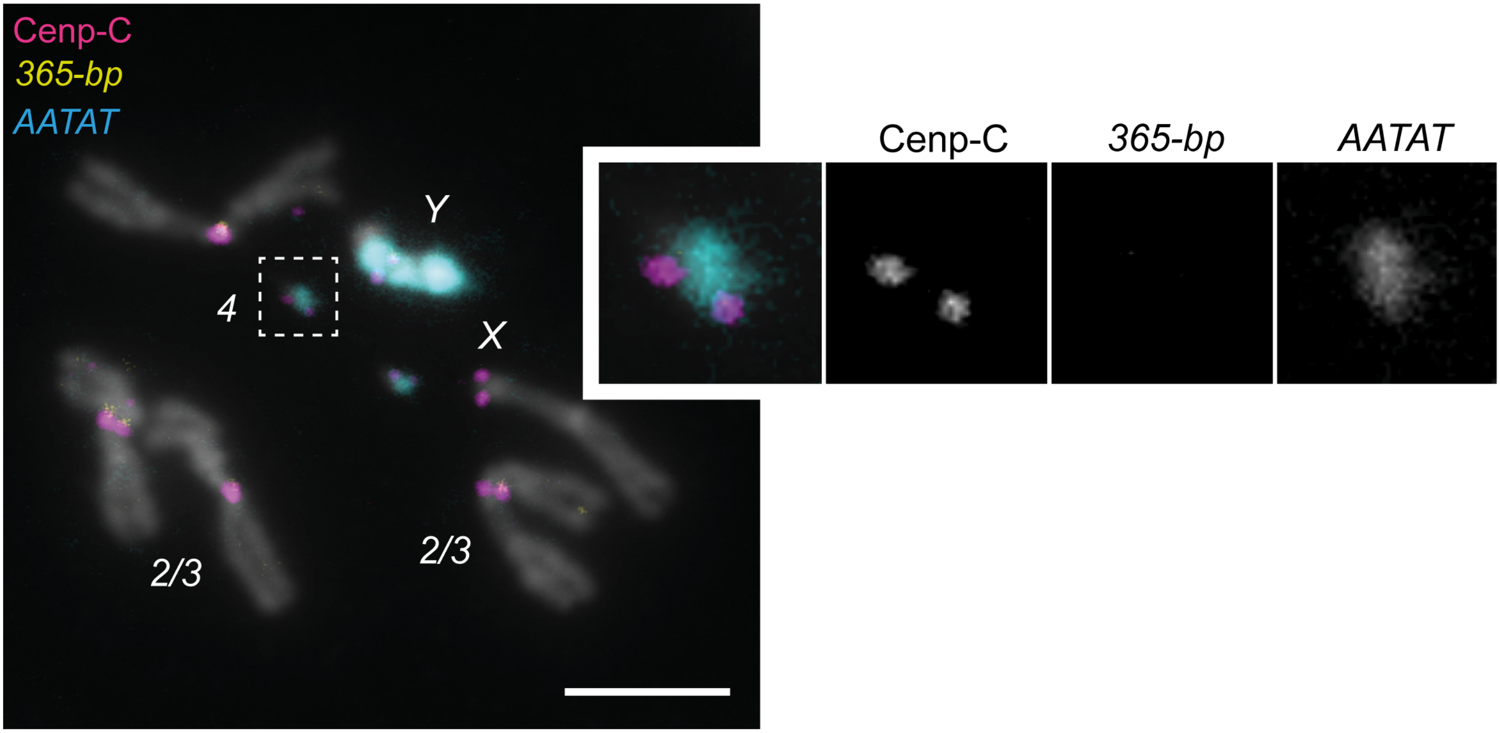

### Sup Fig7

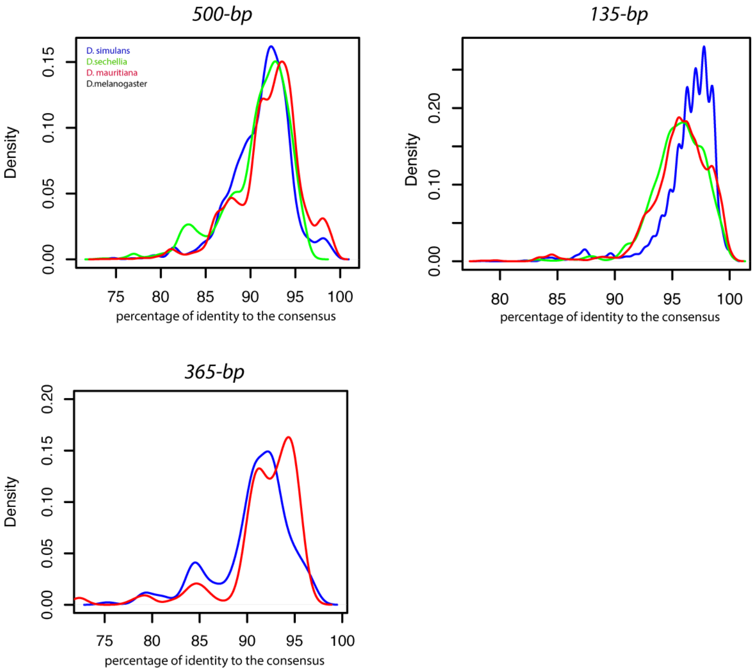

### Sup Fig8

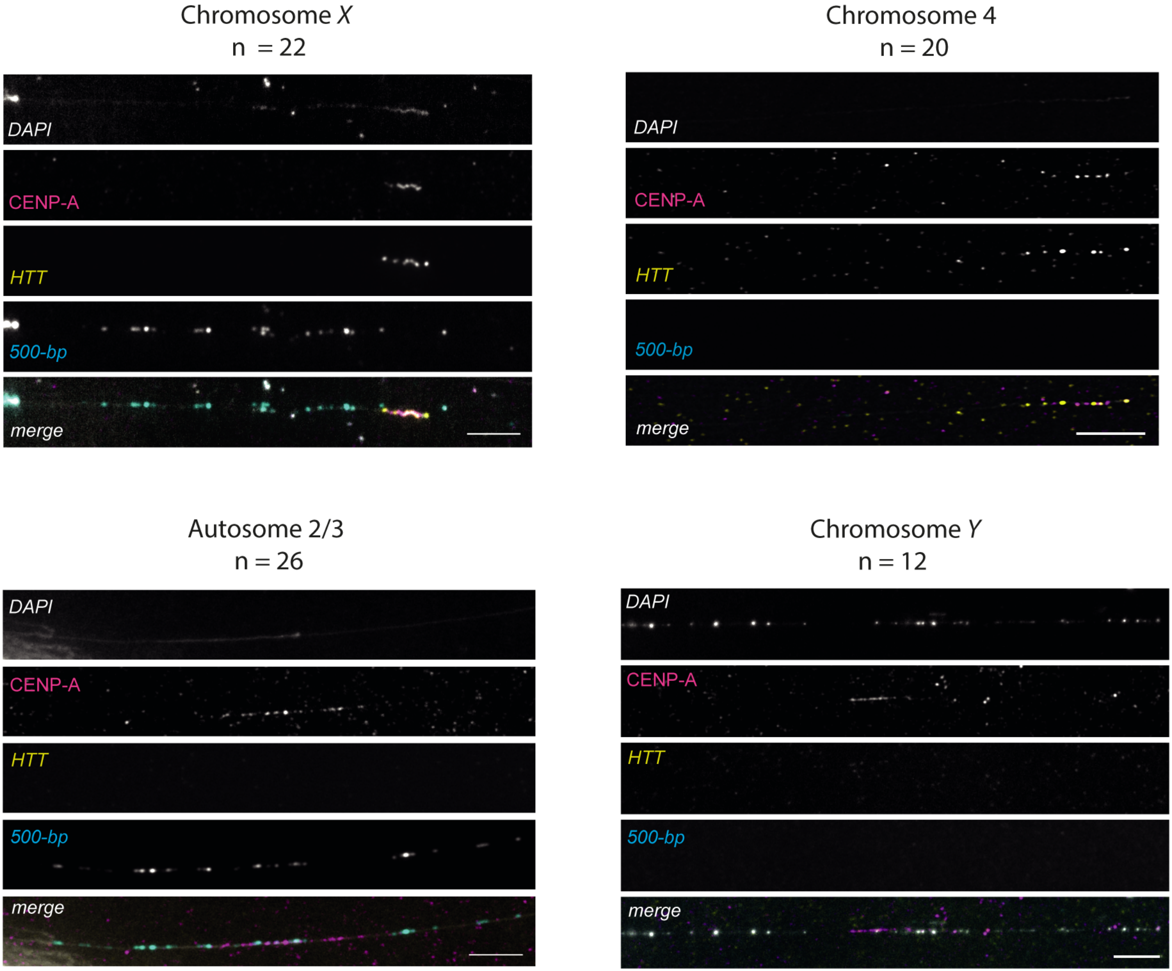

### Sup Fig9

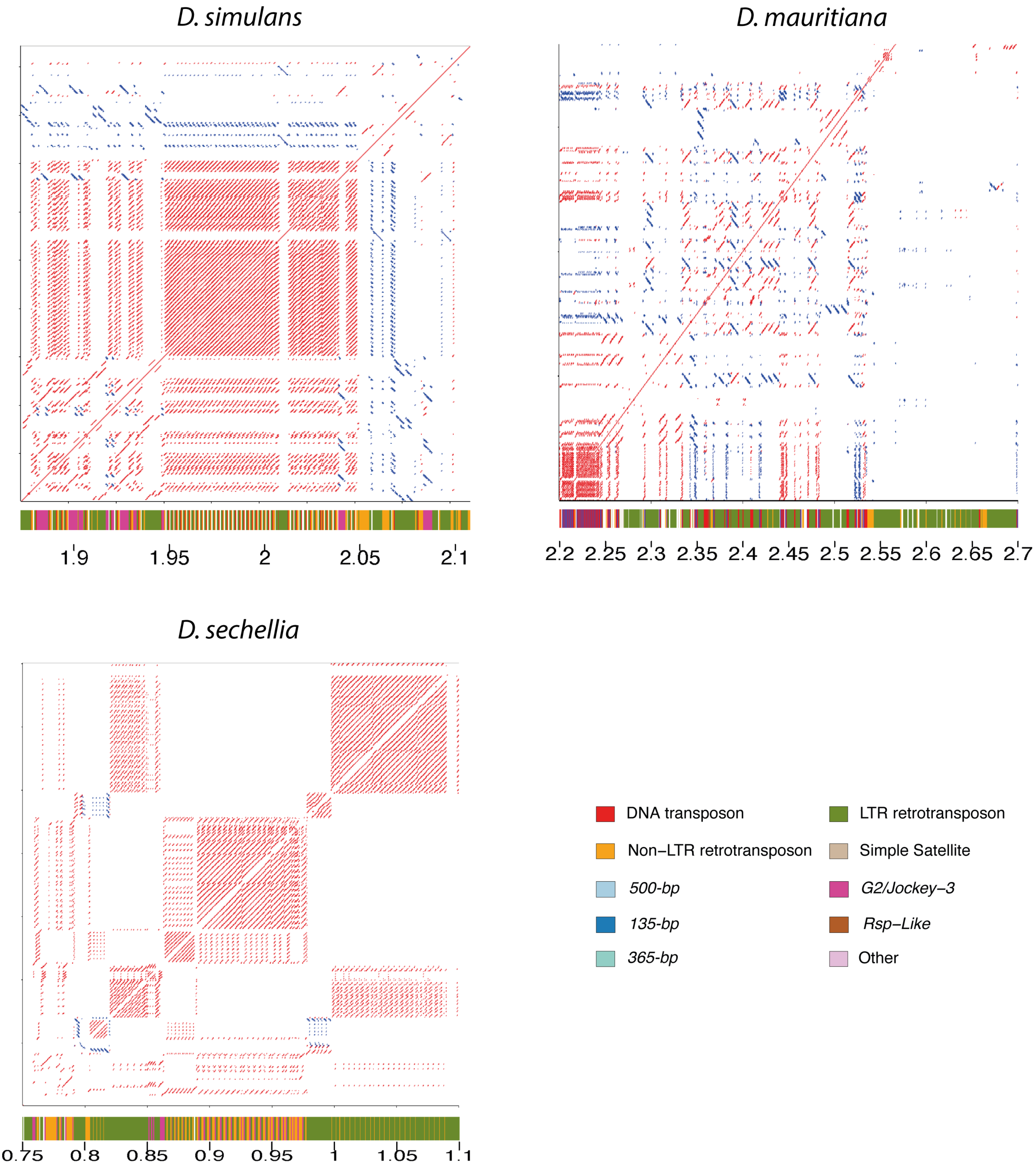

### Sup Fig10

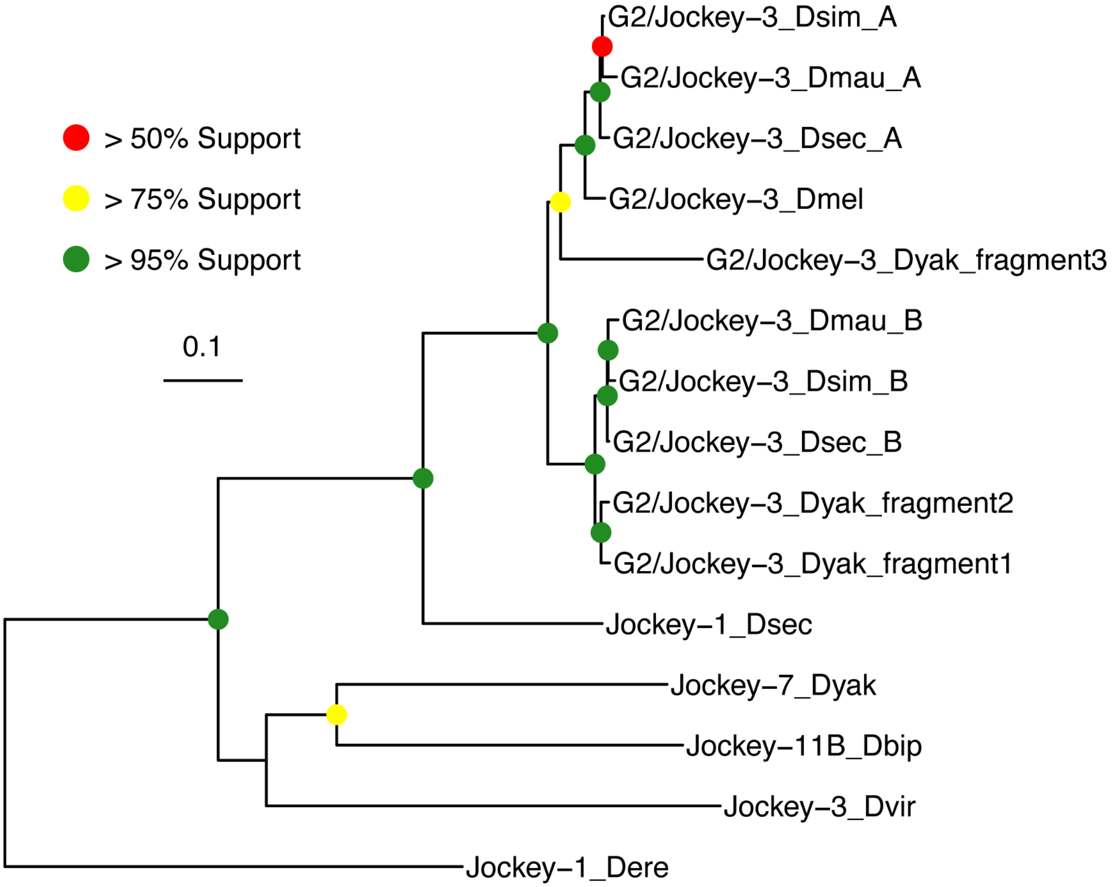

### Sup Fig11

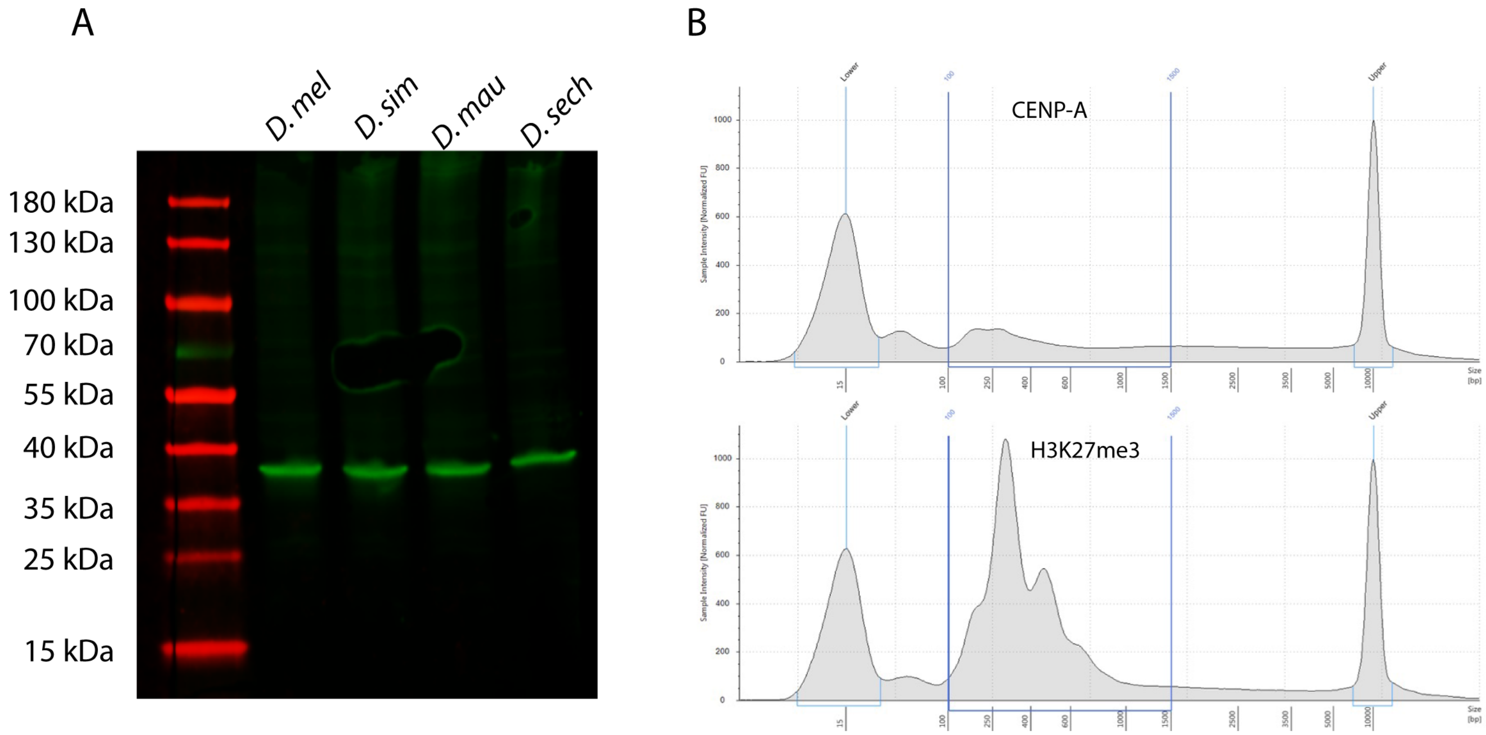
